# Zygomorphic flowers have fewer potential pollinator species

**DOI:** 10.1101/743872

**Authors:** Jeremy B. Yoder, Giancarlo Gomez, Colin J. Carlson

## Abstract

Botanists have long identified bilaterally symmetrical (zygomorphic) flowers with more specialized pollination interactions than radially symmetrical (actinomorphic) flowers. Zygomorphic flowers facilitate more precise contact with pollinators, guide pollinator behaviour, and exclude less effective pollinators. However, whether zygomorphic flowers are actually visited by a smaller subset of available pollinator species has not been broadly evaluated. We compiled 53,609 floral visitation records in 159 communities and classified the plants’ floral symmetry. Globally and within individual communities, plants with zygomorphic flowers are indeed visited by fewer species. At the same time, zygomorphic flowers share a somewhat larger proportion of their visitor species with other co-occurring plants, and have particularly high sharing with co-occurring plants that also have zygomorphic flowers. Visitation sub-networks for zygomorphic species also show differences that may arise from reduced visitor diversity, including greater connectance, greater web asymmetry, and lower coextinction robustness of both plants and visitor species — but these changes do not necessarily translate to whole plant-visitor communities. These results provide context for widely documented associations between zygomorphy and diversification and imply that species with zygomorphic flowers may face greater risk of extinction due to pollinator loss.

## INTRODUCTION

An axiom of pollination ecology is that flowers with bilateral symmetry are more specialized than flowers with radial symmetry [1–4]. “Specialized,” however, has multiple meanings in evolution and ecology, which are not mutually exclusive. Specialization may refer to a derived character state in a phylogenetic context [1,3], or the degree to which a flower manipulates pollinator behaviour [5], or it may refer to association with a particular set of pollinators (i.e., pollination syndromes; [6,7]) — or, finally, it may refer to association with fewer available pollinator species than seen for comparable plant species [8]. Zygomorphic flowers are derived within the angiosperms [9–11], and extensive research examines how floral structure attracts, guides, or excludes pollinators [5,6,12–15]. However, data addressing the fourth sense in which zygomorphic flowers may be specialized — association with fewer pollinators than otherwise expected — are surprisingly sparse.

Floral symmetry has been recognized as an important feature of angiosperm diversity since at least the 18th Century [4]. Modern treatments identified zygomorphy as derived, and hypothesized that zygomorphic forms facilitate more effective pollination [1,3,16]. Zygomorphy is associated with greater diversification rates [17–19], consistent with the hypothesis that using fewer or more constant pollinators creates more opportunities for reproductive isolation [20]. Greater pollination specialization might also interact with global patterns of diversity, such as latitudinal gradients [21,22]: recent syntheses find evidence that biotic interactions are stronger in the tropics [23,24], though assessments of latitudinal effects on pollination specifically have mixed results [25,26].

Floral symmetry has been considered as an element of pollination syndromes [6,7], but to our knowledge, documentation that zygomorphic flowers associate with fewer pollinator species is restricted to anecdotal observations (e.g. [1,3,5]). Broad confirmation of this understanding would illuminate the research linking pollination to diversification [1,16,18,20,27]. Ecologically, the use of fewer pollinators by zygomorphic flowers may have implications for factors ranging from species’ geographic extents to risks of extinction due to pollinator loss.

If floral symmetry creates systematic differences in pollinator associations, these differences should manifest in floral visitation networks. Plant-pollinator associations have been prominent case studies in investigations of ecological networks’ structure and assembly [28–30], geographic variation [25,26], and evolutionary stability [31]. Databases of floral visitation networks have global coverage, recording visitor diversity and sharing among co-occurring angiosperm species across a variety of contexts. Here, we compile a global dataset of floral visitation records to test the hypothesis that zygomorphic flowers have fewer visitor species, and examine how this effect may shape plant-pollinator networks.

## METHODS

We compiled floral visitation networks from the Web of Life (www.web-of-life.es) and Interaction Web DataBase (www.nceas.ucsb.edu/interactionweb). Networks varied widely in size, attributable to heterogeneity in the time periods over which they were studied, their observation protocols, and the geographic ranges of communities represented (Figure 1). We accounted for this in subsequent analyses by simplifying visitation into a binary state (the most common recording mode), and by examining contrasts within individual networks or treating network identity as a random effect (see below).

**Figure 1.**
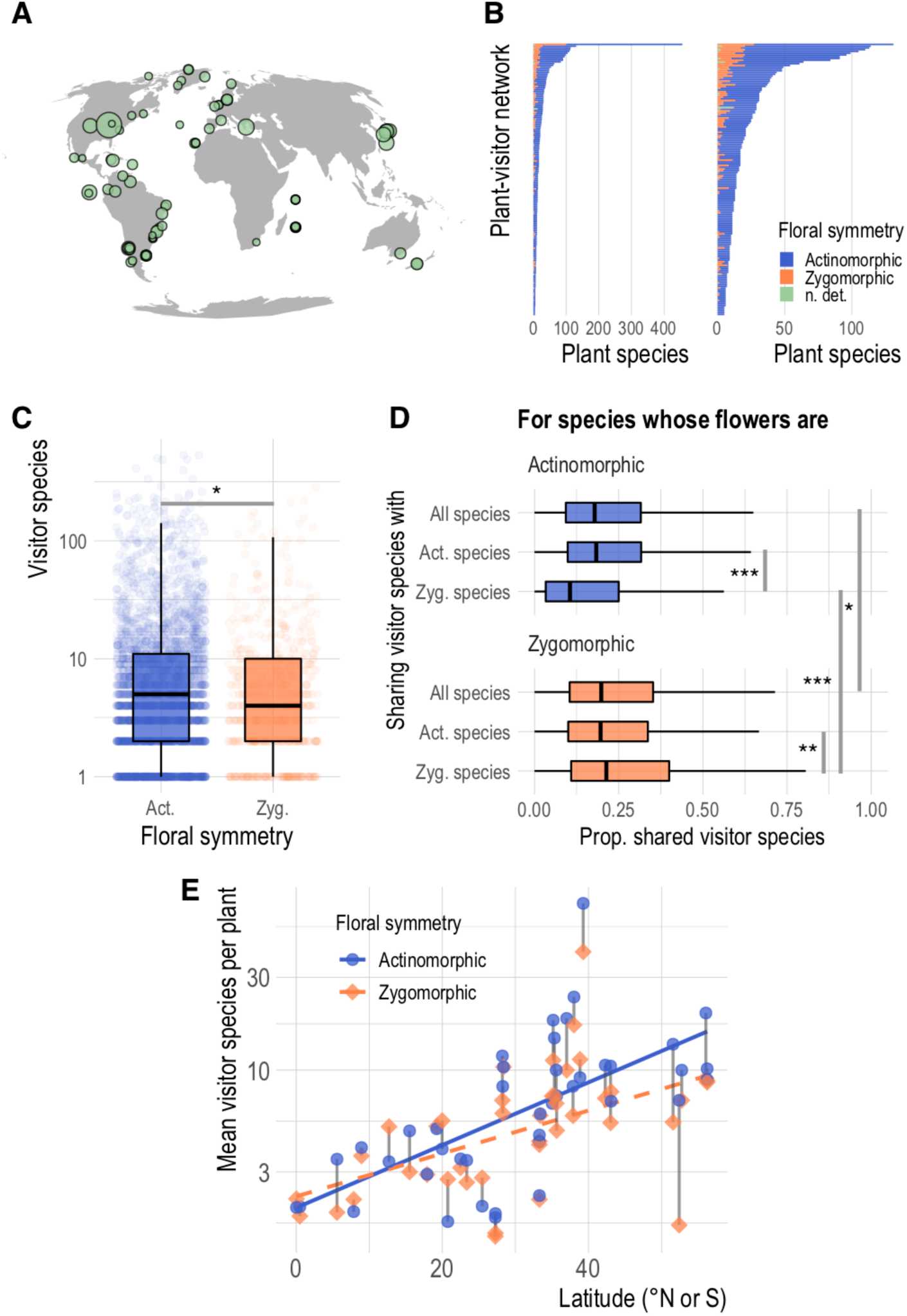
The global plant visitation dataset. (A) Global distribution of the plant-visitor networks, with points scaled to indicate numbers of plant taxa. (B) Bar plots giving the number of plant taxa for each network, coloured according to floral symmetry (left, full dataset, right, all networks except for the largest, to provide better visibility. (C) Visitor species per plant, grouped by floral symmetry. (D) Sharing of visitor species with all co-occurring plant species, and co-occurring species with each type of floral symmetry. (E) Mean visitor species per plant in sub-networks for plants with differing symmetries, with grey lines linking points for zygomorphic and actinomorphic taxa from the same network, plotted against latitude of the network location. In (C) and (D), asterisks indicate significant differences in one-tailed Wilcoxon tests: * p < 0.01, ** p < 10^−4^, *** p < 10^−5^.

We identified unique plant taxa across all networks (hereafter ‘species’; identification of plants and visitors varied in resolution, but 95% of records had plants identified to species), and classified their floral symmetry based on taxonomic knowledge, formal descriptions of species or higher taxa, and, when necessary, inspecting images of herbarium sheets or reliably identified fresh flowers. We determined symmetry based on the perianth, but in ambiguous cases also considered symmetry of the androecium or gynoecium; for example, a flower with a very nearly radially symmetric corolla, but stamens arrayed in a bilaterally symmetric manner, would be coded as zygomorphic. In some cases, we classified symmetry not based on individual flowers but on flowering heads (e.g., we considered species in the Asteraceae actinomorphic). We removed species from the working dataset if we were unable to find authoritative descriptions or images, or if they were wind-pollinated (“n. det.”, Figure 1B). Data and scripts are posted to Dryad, at https://datadryad.org/10.5061/dryad.gxd2547j3

We conducted analysis in R v. 4.0 [32]. For each plant species in the dataset, we totalled the visitor species recorded, and calculated an index of visitor species sharing, the proportion of visitor species to each plant species that also visit each other co-occurring plant species, averaged across the co-occurring plant species. We calculated sharing with all plants in the same network, and sharing with plants in the same network having each type of floral symmetry. We examined the structure of complete networks, and of the sub-networks of visitors to plants with each type of floral symmetry in each community, calculating connectance (the realized proportion of possible plant-visitor links [33]), web asymmetry (the degree to which visitor species outnumber plant species or vice-versa), and coextinction curves (the relationship between species losses in one trophic level and species losses in another [34]) using the `networklevel()` and `second.extinct()` functions in the `bipartite` package [35].

To test for phylogenetic signal in floral symmetry, visitor species count, and visitor species sharing, and to control for phylogeny in subsequent analysis, we mapped taxa in our dataset to a recently published time-calibrated supertree of the seed plants (the “ALLMB” supertree of [36]), using the `congeneric.merge()` function from the package `pez` [37] to add species to the tree if they were not already included. We tested for phylogenetic signal using the `phylosignal` package [38], estimating Blomberg’s K and K* [39], Abouheif’s C_mean_ [40], Moran’s I [41], and Pagel’s λ [42] statistics. We performed principal component analysis of the phylogenetic distance matrix for all plant taxa and used phylogenetic distance principal components as covariates in models fitted to explain variation in visitor count and sharing.

We tested the hypotheses that visitor species count and visitor species sharing differed with respect to floral symmetry by fitting Bayesian multilevel regression models using the brms package [43,44]. Competing models explained visitor count and sharing with a group effect (analogous to a random effect in a ML framework) of source network identity and possible population (i.e., fixed) effects of floral symmetry, latitude, and the first two principal components of phylogenetic distance (which jointly explained 61% of variation). We compared model fit in terms of expected log pointwise predictive density (ELPD) using leave-one-out cross-validation, implemented in `brms` with the `LOO()` function [45].

## RESULTS

We compiled 159 networks, recording 53,609 observed visits to 2,700 angiosperm species (Figure 1A; Supplementary Table 1). We were able to classify floral symmetry for 2,685 plant species, and were able to place 2,582 of these in the time-calibrated supertree [36]. Globally, and in individual networks, zygomorphic flowers were a minority: 498 species (18%) were zygomorphic; only five networks had more zygomorphic than actinomorphic species, while 70 lacked any (Figure 1B). Globally, the number of visitor species to zygomorphic flowers was significantly smaller than that for actinomorphic flowers (median 4 pollinators per zygomorphic species with zygomorphic flowers versus 5 per actinomorphic species; p = 0.003, one-tailed Wilcoxon test; Figure 1C). We found significant phylogenetic signal for floral symmetry (C_mean_ = 0.85, Moran’s I = 0.10, K = 0.20, K* = 0.22, and λ = 0.89; p < 0.001 for all); visitor species count deviated from the null models for C_mean_, I, and λ (p < 0.001 for each) but not for K and K*.

At the same time, zygomorphic flowers shared a higher proportion of their visitor species with co-occurring plants (median 0.20 for zygomorphic versus 0.18 for actinomorphic, p = 0.003; Figure 1D). Plants also had greater sharing with co-occurring plants of the same floral symmetry (actinomorphic species, median sharing of 0.24 with actinomorphic species versus 0.19 with zygomorphic species, p = 0.003; zygomorphic species, median sharing of 0.30 with zygomorphic species versus 0.27 with actinomorphic species; p < 10^−5^; Figure 1D).

Thirty-nine networks comprising 46,026 visitation records included enough plants of each symmetry type (at least five) to compare sub-networks based on symmetry — that is, to compare networks for plants with different symmetry having access to the same pool of possible visitor species. Zygomorphic sub-networks had significantly fewer mean visitor species per plant (one-tailed paired Wilcoxon test, p < 0.001; Figure 2A), but only marginally greater visitor sharing (p = 0.06). Zygomorphic sub-networks had significantly greater connectance (p < 10^−4^), greater asymmetry (p < 0.001), and lower coextinction robustness, as measured by the exponent of the coextinction curve, for both plants (p < 10^−5^) and visitors (p < 10^−5^). Among 89 complete networks that included at least one zygomorphic plant, a higher proportion of zygomorphic plants was associated with greater visitor sharing (Figure 2B; product-moment correlation on log-transformed values = 0.22, p = 0.04) and greater connectance (cor = 0.21, p = 0.04), paralleling the sub-network patterns. However, networks with more zygomorphic flowers also had lower asymmetry (cor = -0.26, p = 0.01), and greater visitor coextinction robustness (cor = 0.27, p = 0.01), the opposite of patterns in symmetry-based sub-networks. The proportion of zygomorphic flowers was not correlated with the mean number of visitors per plant or plant coextinction robustness.

**Figure 2.**
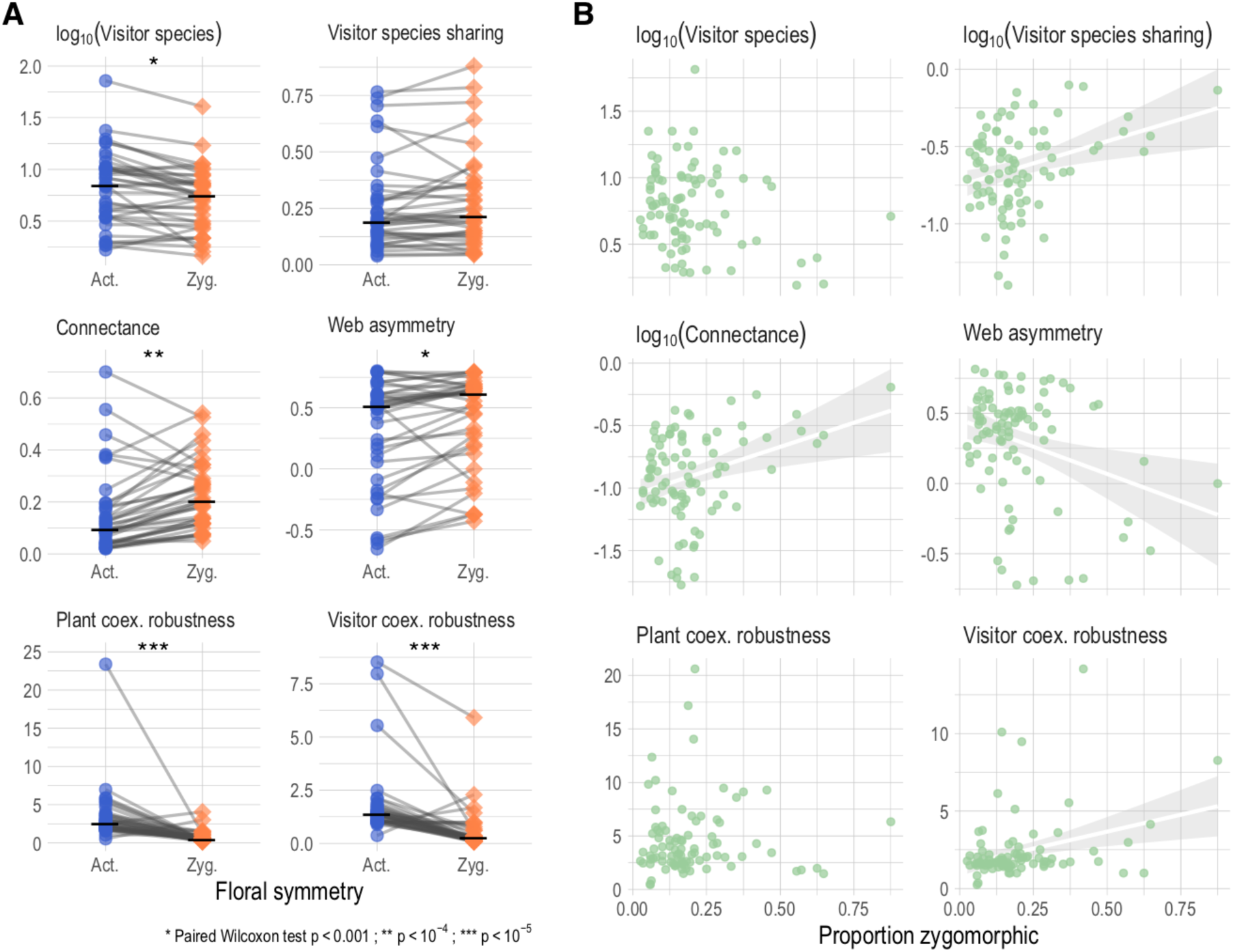
Floral symmetry and plant-visitor network structure. (A) Network descriptive statistics for sub-networks based on floral symmetry. Grey lines link points representing zygomorphic and actinomorphic sub-networks from the same source network, horizontal black bars mark the median value within each floral symmetry type, and asterisks mark significant differences in one-tailed paired Wilcoxon tests. (B) Scatterplots of relationships between the proportion of zygomorphic flowers in a network and the set of network structure metrics in A; linear regression lines (white with grey 95% CI) are given when the correlation is significant with p ≤ 0.05.

The best-fit model explaining visitor count included floral symmetry, latitude, an interaction between symmetry and latitude, and phylogeny in addition to the group effect of network identity (R^2^ = 0.28; ΔELPD ≥ 3.8 versus all other models; Supplementary Tables 2, 3). For visitor sharing, the best-fit model also included all terms; but all simpler models that included an effect of symmetry had ΔELPD ≤ 0.6 (R^2^ = 0.66 for all such models; Supplementary Tables 4, 5).

## DISCUSSION

Zygomorphic flowers have long been considered to be more specialized, which could mean that they are visited by fewer pollinator species. We find that, globally and at the level of individual communities, plants with zygomorphic flowers do indeed have fewer visitor species, and that the visitation networks of plants with zygomorphic symmetry may be structured by this difference (Figure 1C,D; Figure 2A). Sub-networks of plants with zygomorphic flowers show greater connectance, greater asymmetry, and lower coextinction robustness for both plants and visitors (Figure 2A); however, these patterns do not necessarily translate to whole networks (Figure 2B). Visitor species count is correlated with latitude north or south (Figure 1E), and both floral symmetry and visitor count show significant phylogenetic structure. Bayesian multilevel regressions accounting for these confounding effects nevertheless find significant effects of floral symmetry on visitor species count and sharing.

The visitation records we examine have limitations for assessing pollination specialization. Many of the original studies do not evaluate visitors’ pollen transfer, and often record visits as a binary, while in reality floral visitors vary considerably in visitation frequency and effectiveness as pollinators. Thus, the effective pollinators of the plants in our dataset are likely a subset of recorded visitors. However, we think it unlikely that data restricted to effective pollinators would reverse the qualitative patterns we see, because a reversal would require systematic bias based on floral symmetry in recording visits.

Our findings that zygomorphic flowers share more visitor species make sense given the other aspects of pollinator specialization associated with zygomorphy. Arithmetically, a plant with fewer visitors can more easily share a high proportion of them even if the absolute number of visitors shared is low, but higher sharing by zygomorphic species probably also reflects network structure. Plant-visitation networks are generally nested [30], so the fact that zygomorphic species tend to have fewer visitor species means that their visitors are more likely to interact with many other plant taxa. Zygomorphic flowers, which often manipulate pollinator behaviour, apply pollen to specific parts of pollinators’ bodies, or attract pollinators that show greater constancy over a single foraging trip, may be better able to tolerate elevated risk of receiving heterospecific pollen due to higher visitor sharing [1,20]. The fact that the highest rate of sharing we see in our data is between zygomorphic flowers and other co-occurring zygomorphic species is consistent with this hypothesis.

In our compiled dataset, plants with both types of floral symmetry had more visitor species in communities farther from the equator (Figure 1E). This could reflect bias in the resolution of species in more tropical communities, as there are more undescribed species in the tropics, and in the Global South [46]. Networks farther from the equator do have more plant and visitor species recorded (cor = 0.54 for plants, cor = 0.65 for visitors, p < 10^−5^ for both; see Figure 1A). However, the correlation weakens or disappears if the comparison is with latitude rather than distance from the equator (i.e., treating distance south of the equator as different from distance north: cor = 0.3, p = 0.69 for plants; cor = 0.16, p = 0.05 for visitors), suggesting this is not simply an issue of better taxonomy in the Global North. Moreover, if plant and visitor species numbers show similar correlations with distance from the equator, we might expect that the number of visitors *per* plant would be constant across latitudes. An alternative explanation is that floral visitation is more specialized in the tropics, independent of floral symmetry. Testing this hypothesis with greater rigor is beyond the scope of our data, but it would be consistent with recent syntheses finding stronger effects of biological interactions in the tropics [23,24].

An ecological association between floral symmetry and pollinator diversity may explain evolutionary associations, across the angiosperms, between floral zygomorphy and diversification [17,18,47]. The direction of the causal relationship, however, remains ambiguous. It may be that association with fewer pollinators creates more opportunities to evolve reproductive isolation [1,20,47] or increases the number of plant species that can be supported by a given community of pollinators [18]; or it may be that more specialized pollination evolves in response to greater diversity as co-occurring species must subdivide available pollinators more finely [27].

Finally, our results have important implications for conservation. We find lower coextinction robustness for sub-networks of plants with zygomorphic flowers, which may be explained by higher connectance and asymmetry of these sub-networks (Figure 2A) and higher visitor sharing among zygomorphic flowers (Figure 1D). These patterns are not necessarily borne out at the level of whole communities (Figure 2B), suggesting that larger communities may be robust to disturbances that endanger their most specialized members. Nevertheless, our analysis does imply that plant taxa with zygomorphic flowers are at greater risk of extinction due to pollinator loss. Despite significant uncertainty in the magnitude of losses, pollinators are widely known to be in rapid decline due to pesticide use, habitat degradation, and emerging infections [48,49]. The patterns we find are coarse, but simple rubrics for triage are critical for conserving the more than 300,000 species of angiosperms, most of which will never benefit from individualized conservation assessment. Perhaps more importantly, our results support the idea that “compartments” of the global plant-pollinator network must be targets for holistic conservation, focused on preserving interactions and functionality where the network is most fragile [50].

## ACKNOWLEDGEMENTS

JBY was supported by start-up funds from California State University Northridge, and CJC by a postdoctoral fellowship from the Georgetown Environment Initiative. We thank Shweta Bansal for helpful conversations and Stephen Sondheim for teaching us that a faboid legume can begin an adventure.

## DATA, CODE, AND MATERIALS

Data including floral symmetry annotations, visitor counts and sharing, and network structure statistics are available with scripts on Dryad, https://datadryad.org/10.5061/dryad.gxd2547j3

## AUTHORS’ CONTRIBUTIONS

JBY and CJC conceived and designed the study; GG annotated floral symmetry with supervision from JBY, and JBY and GG conducted analysis with code provided by CJC; JBY drafted the paper in consultation with CJC and GG. All gave final approval for publication and agree to be held accountable for the work performed.

