## Supplementary Information for "Zygomorphic flowers have fewer potential pollinator species" (from another version of this preprint: version 5, 2020-07-03)

for

### Zygomorphic flowers have fewer visitors

Last saved by Jeremy Yoder, 30 April 2020

**Supplementary Table 1.** Data sources for plant visitation observations.

| Network identifier | Source[1] | Latitude | Reference |
|---|---|---|---|
| M_PL_064 | WoL | -20.75 | Abreu, C.R.M. & Vieira, M.F. (2004) Lundiana, 5, 129,Äì134. |
| M_PL_065 | WoL | 19.5 | Arizmendi, M.C. & Ornelas, J.F. (1990) Biotropica, 22, 172,Äì180. |
| M_PL_001 | WoL | -33.283334 | Arroyo, M.T.K., R. Primack & J.J. Armesto. 1982. Amer. J. Bot. 69:82-97. |
| M_PL_002 | WoL | -33.283334 | |
| M_PL_003 | WoL | -33.283334 | |
| M_PL_004 | WoL | 46.553731 | Barrett, S. C. H., and K. Helenurm. 1987. Canadian Journal of Botany 65:2036-2046. |
| M_PL_058 | WoL | 42.2973 | Bartomeus, I., Vilà, M. & Santamaria, L., 2008. Oecologia 155: 761-770. |
| M_PL_049 | WoL | 56.066667 | Bek S (2006). MSc thesis (Univ of Aarhus, Aarhus, Denmark). |
| M_PL_059 | WoL | -8.505 | Bezerra ELS, Machado ICS, Mello MAR. 2009. Journal of Animal Ecology 78:1096-1101. |
| M_PL_046 | WoL | 56.066667 | Bundgaard, M. 2003. University of Aarhus. Aarhus, Denmark. |
| M_PL_066 | WoL | -22.5 | Canela, M.B.F. 2006. Ph.D thesis. Universidade Estadual de Campinas, Brazil. |
| M_PL_005 | WoL | 38.841784 | Clements, R. E., and F. L. Long. 1923, Washington, D.C., USA, Carnegie Institute of Washington. |
| M_PL_006 | WoL | 52.762395 | Dicks, LV, Corbet, SA and Pywell, RF 2002. J. Anim. Ecol. 71: 32-43. |
| M_PL_007 | WoL | 52.413173 | |
| M_PL_047 | WoL | 56.066667 | Dupont & Olesen 2009. J. Animal Ecology, 78:346-353. |
| M_PL_048 | WoL | 56.1 | |
| M_PL_008 | WoL | 28.216667 | Dupont YL, Hansen DM and Olesen JM 2003. Ecography 26:301-310. |
| M_PL_010 | WoL | 74.5 | Elberling & Olesen unpubl. |
| M_PL_009 | WoL | 68.35 | Elberling, H. & Olesen, J. M. 1999. Ecography 22:314-323. |
| M_PL_068 | WoL | 1.25 | Gutierrez Zamora, E.A. & Rojas Nossa, S.V. 2001. BSc. Thesis. Universidad Nacional de Colombia, Colombia. |
| M_PL_016 | WoL | 37.016667 | Herrera, J. 1988. Journal of Ecology 76: 274-287. |
| M_PL_014 | WoL | 81.816667 | Hocking, B. 1968. Oikos 19:359-388. |
| M_PL_041 | WoL | 15.518889 | Ingversen TT 2006. MSc thesis (Univ of Aarhus, Aarhus, Denmark). |
| M_PL_040 | WoL | 18.353889 | |

| Network identifier | Source1 | Latitude | Reference |
|---|---|---|---|
| M_PL_057 | WoL | 35.166667 | Inoue et al 1990. Contribution from the Biological Laboratory, Kyoto University Univ. 27, 377-463. |
| M_PL_019 | WoL | -36.45 | Inouye, D. W., and G. H. Pyke. 1988. Australian Journal of Ecology 13:191-210. |
| M_PL_061_01 | WoL | -4.666667 | Kaiser-Bunbury CN, Vázquez DP, Stang M, Ghazoul J. 2014. Ecology, 95: 3314-3324. |
| M_PL_061_02 | WoL | -4.666667 | |
| M_PL_061_03 | WoL | -4.666667 | |
| M_PL_061_04 | WoL | -4.666667 | |
| M_PL_061_05 | WoL | -4.666667 | |
| M_PL_061_06 | WoL | -4.666667 | |
| M_PL_061_07 | WoL | -4.666667 | |
| M_PL_061_08 | WoL | -4.666667 | |
| M_PL_061_09 | WoL | -4.666667 | |
| M_PL_061_10 | WoL | -4.666667 | |
| M_PL_061_11 | WoL | -4.666667 | |
| M_PL_061_12 | WoL | -4.666667 | |
| M_PL_061_13 | WoL | -4.666667 | |
| M_PL_061_14 | WoL | -4.666667 | |
| M_PL_061_15 | WoL | -4.666667 | |
| M_PL_061_16 | WoL | -4.666667 | |
| M_PL_061_17 | WoL | -4.666667 | |
| M_PL_061_18 | WoL | -4.666667 | |
| M_PL_061_19 | WoL | -4.666667 | |
| M_PL_061_20 | WoL | -4.666667 | |
| M_PL_061_21 | WoL | -4.666667 | |
| M_PL_061_22 | WoL | -4.666667 | |
| M_PL_061_23 | WoL | -4.666667 | |
| M_PL_061_24 | WoL | -4.666667 | |
| M_PL_061_25 | WoL | -4.666667 | |
| M_PL_061_26 | WoL | -4.666667 | |
| M_PL_061_27 | WoL | -4.666667 | |
| M_PL_061_28 | WoL | -4.666667 | |
| M_PL_061_29 | WoL | -4.666667 | |
| M_PL_061_30 | WoL | -4.666667 | |
| M_PL_061_31 | WoL | -4.666667 | |
| M_PL_061_32 | WoL | -4.666667 | |

| Network identifier | Source[1] | Latitude | Reference |
|---|---|---|---|
| M_PL_061_33 | WoL | -4.666667 | |
| M_PL_061_34 | WoL | -4.666667 | |
| M_PL_061_35 | WoL | -4.666667 | |
| M_PL_061_36 | WoL | -4.666667 | |
| M_PL_061_37 | WoL | -4.666667 | |
| M_PL_061_38 | WoL | -4.666667 | |
| M_PL_061_39 | WoL | -4.666667 | |
| M_PL_061_40 | WoL | -4.666667 | |
| M_PL_061_41 | WoL | -4.666667 | |
| M_PL_061_42 | WoL | -4.666667 | |
| M_PL_061_43 | WoL | -4.666667 | |
| M_PL_061_44 | WoL | -4.666667 | |
| M_PL_061_45 | WoL | -4.666667 | |
| M_PL_061_46 | WoL | -4.666667 | |
| M_PL_061_47 | WoL | -4.666667 | |
| M_PL_061_48 | WoL | -4.666667 | |
| M_PL_060_01 | WoL | -20.7 | Kaiser-Bunbury, C. N., S. Muff, J. Memmott, C. B. Müller, and A. Caflisch. 2010. Ecology Letters 13:442-452. |
| M_PL_060_02 | WoL | -20.7 | |
| M_PL_060_03 | WoL | -20.7 | |
| M_PL_060_04 | WoL | -20.7 | |
| M_PL_060_05 | WoL | -20.7 | |
| M_PL_060_06 | WoL | -20.7 | |
| M_PL_060_07 | WoL | -20.7 | |
| M_PL_060_08 | WoL | -20.7 | |
| M_PL_060_09 | WoL | -20.7 | |
| M_PL_060_10 | WoL | -20.7 | |
| M_PL_060_11 | WoL | -20.7 | |
| M_PL_060_12 | WoL | -20.7 | |
| M_PL_060_13 | WoL | -20.7 | |
| M_PL_060_14 | WoL | -20.7 | |
| M_PL_060_15 | WoL | -20.7 | |
| M_PL_060_16 | WoL | -20.7 | |
| M_PL_060_17 | WoL | -20.7 | |
| M_PL_060_18 | WoL | -20.7 | |
| M_PL_060_19 | WoL | -20.7 | |

| Network identifier | Source1 | Latitude | Reference |
|---|---|---|---|
| M_PL_060_20 | WoL | -20.7 | |
| M_PL_060_21 | WoL | -20.7 | |
| M_PL_060_22 | WoL | -20.7 | |
| M_PL_060_23 | WoL | -20.7 | |
| M_PL_060_24 | WoL | -20.7 | |
| M_PL_054 | WoL | 35.033333 | Kakutani, T., T. Inoue, M. Kato and H. Ichihashi 1990. Contribution from the Biological Laboratory, Kyoto University, 27: 465-521. |
| M_PL_055 | WoL | 35.65 | Kato & Miura 1996. Kyoto University, Vol. 29: 1-48 |
| M_PL_056 | WoL | 35.583333 | Kato et al. 1993. Contributions from the Biological Laboratory, Kyoto University 28: 119-172 |
| M_PL_044 | WoL | 28.377248 | Kato M 2000. Contributions from the Biological Laboratory, Kyoto University 29:157-252. |
| M_PL_021 | WoL | 35.333333 | Kato, M., Kakutani, T., Inoue, T. and Itino, T. 1990. Contributions from the Biological Laboratory, Kyoto University, 27, 309-375. |
| M_PL_020 | WoL | 81.816667 | Kevan P. G. 1970. Ph.D. thesis, University of Alberta, Edmonton, 399 pp. |
| M_PL_069_01 | WoL | -27.27 | Kohler, G. 2011. MSc. Thesis. Universidade Federal do Paraná, Brazil. |
| M_PL_069_02 | WoL | -27.26 | |
| M_PL_069_03 | WoL | -27.26 | |
| M_PL_070 | WoL | 19.23 | Lara, C. 2006. Ecoscience, 13, 23,Äì29. |
| M_PL_067 | WoL | -7.87 | Las Casas, F.M.G., Azevedo Júnior, S.M. & Dias Filho, M.M. 2012 Brazilian Journal of Biology, 72, 51-58. |
| M_PL_045 | WoL | 71 | Lundgren R, Olesen JM 2005. Arc Antarc Alp Res 37:514-520. |
| M_PL_026 | WoL | -0.5 | McMullen, C. K. 1993. Pan-Pacific Entomologist 69:95-106 |
| M_PL_022 | WoL | -34.166667 | Medan, D., N. H. Montaldo, M. Devoto, A. Mantese, V. Vasellati, and N. H. Bartoloni. 2002. Arctic Antarctic and Alpine Research 34:233-241. |
| M_PL_023 | WoL | -33 | |
| M_PL_017 | WoL | 51.574994 | Memmott J. 1999. Ecology Letters 2:276-280. |
| M_PL_037 | WoL | 56.248737 | Montero AC 2005. MSc thesis (Univ of Aarhus, Aarhus, Denmark). |
| M_PL_038 | WoL | 56.238737 | |
| M_PL_043 | WoL | 56.238737 | |
| M_PL_024 | WoL | 75 | Mosquin, T., and J. E. H. Martin. 1967. Canadian Field Naturalist 81:201-205. |

| Network identifier | Source1 | Latitude | Reference |
|---|---|---|---|
| M_PL_025 | WoL | 36.075391 | Motten, A. F. 1982. Ecological Monographs 56:21-42. |
| M_PL_012 | WoL | 28.127842 | Olesen unpubl. |
| M_PL_018 | WoL | 56.238737 | |
| M_PL_036 | WoL | 39.466667 | |
| M_PL_011 | WoL | -20.348404 | Olesen, J.M., Eskildsen, L.I. & Venkatasamy, S. 2002. Div. Distr., 8:181-192. |
| M_PL_013 | WoL | -29.616667 | Ollerton, J., S. D. Johnson, L. Cranmer, and S. Kellie. 2003. Annals of Botany 92:807-834. |
| M_PL_035 | WoL | 17.916667 | Percival, M. 1974. Biotropica, 6, 104-129. |
| M_PL_015 | WoL | 38.014466 | Petanidou, T. 1991. Unp. PhD. Thesis, Aristotelian University, Thessaloniki. |
| M_PL_042 | WoL | -0.949194 | Philipp M, Böcher J, Siegismund HR, Nielsen LR 2006. Ecography 29:531-540. |
| M_PL_027 | WoL | -42.95 | Primack, R.B. 1983. New Zealand J. Bot. 21, 317-333. AP. |
| M_PL_028 | WoL | -43.02823 | |
| M_PL_029 | WoL | -43.099531 | |
| M_PL_031 | WoL | 5.583333 | Ramirez, N. 1989. Biotropica 21, 319-330. |
| M_PL_030 | WoL | 8.933333 | Ramirez, N., and Y. Brito. 1992. Botanical Journal of the Linnean Society 110:277-302. |
| Robertson-1928 | WoL | 39.278958 | Robertson, C. 1929. Flowers and insects: lists of visitors to four hundred and fifty-three flowers. Carlinville, IL, USA, C. Robertson. |
| M_PL_071 | WoL | 0.04 | Rosero, L. 2003. Ph.D. Thesis. Universidade Estadual de Campinas, Brazil |
| M_PL_072_01 | WoL | -37.842015 | Sabatino M., Maceira N., Aizen A. M. 2010. Ecological Applications, 20(6), pp. 1491-1497 |
| M_PL_072_02 | WoL | -37.739035 | |
| M_PL_072_03 | WoL | -37.757207 | |
| M_PL_072_04 | WoL | -37.885354 | |
| M_PL_072_05 | WoL | -37.738483 | |
| M_PL_032 | WoL | 40.133333 | Schemske, D. W., M. F. Willson, M. N. Melampy, L. J. Miller, L. Verner, K. M. Schemske, and L. B. Best. 1978. Ecology 59:351-366. |
| M_PL_033 | WoL | 45.4 | Small, E. 1976. Canadian Field Naturalist 90:22-28. |
| M_PL_034 | WoL | -42 | Smith-Ramírez C., P. Martinez, M. Nuñez, C. González and J. J. Armesto (2005) Botanical Journal of the Linnean Society, 2005, 147, 399-416. |

| Network identifier | Source[1] | Latitude | Reference |
|---|---|---|---|
| M_PL_039 | WoL | 28.268611 | Stald L 2003. Msc thesis (Univ of Aarhus, Aarhus, Denmark). |
| M_PL_050 | WoL | 28.268611 | Stald, L., Valido, A. & Olesen, J. M. 2003. MSc-thesis, Univ. of Aarhus, Denmark. |
| M_PL_051 | WoL | -41.077069 | Vázquez DP. 2002. Ph.D. Dissertation. Department of Ecology and Evolutionary Biology, University of Tennessee, Knoxville. |
| M_PL_063 | WoL | -23.341667 | Vizentin-Bugoni, J., P. K. Maruyama, V. J. Debastiani, L. S. Duarte, B. Dalsgaard & M. Sazima 2016. Journal of Animal Ecology 85: 262-272. |
| M_PL_052 | WoL | 66.966667 | Witt P 1998. BSc thesis. Univ of Aarhus, Aarhus, Denmark. |
| M_PL_053 | WoL | 33.4 | Yamazaki, K. & Kato, M. 2003. Contributions from the Biological Laboratory, Kyoto University 29: 255-318. |
| kaehler_et_al_2005 | IWDB | -25.4583 | Kaehler M., Varassin I.G. & Goldenberg R. 2005. Revista Brasileira de Botânica, 28: 219-228. |
| Kaiser-Bunbury-2017_Bernica | IWDB | -4.6667 | Kaiser-Bunbury CN, Mougal J, Whittington A, Valentin T, Gabriel R, Olesen JM, Blüthgen N. 2017. Nature. DOI: 10.1038/nature21071 |
| Kaiser-Bunbury-2017_Casse-Dent | IWDB | -4.6667 | |
| Kaiser-Bunbury-2017_Copolia | IWDB | -4.6667 | |
| Kaiser-Bunbury-2017_Reserve--La | IWDB | -4.6667 | |
| Kaiser-Bunbury-2017_Rosebelle | IWDB | -4.6667 | |
| Kaiser-Bunbury-2017_Salazie | IWDB | -4.6667 | |
| Kaiser-Bunbury-2017_Tea-Plantation | IWDB | -4.6667 | |
| Kaiser-Bunbury-2017_Trois-Freres | IWDB | -4.6667 | |
| santos-et-al-2010 | IWDB | -12.7 | Santos GMM, Aguiar CML, Mello MAR. 2010. Apidologie, DOI: 10.1051/apido/2009081. |
| varassin_sazima_2012 | IWDB | -19.97638 | Varassin I.G. & Sazima M. 2012. Acta Oecologica, 43: 104-112. |

[1] Sources are either IWDB, the Interaction Web DataBase (www.nceas.ucsb.edu/interactionweb), or WOL, the Web of Life repository (www.web-of-life.es)

**Supplementary Table 2.** Fit (expected likelihood predictive density, ELPD) and variation explained ($R_2$) for Bayesian multilevel models explaining variation in floral visitor count, $N_{visitors}$

| Model* | ELPD ± SE | ΔELPD ± SE | $R_2$ ± SE |
|---|---:|---:|---:|
| (N) | -8649.9 ± 82.1 | 47.1 ± 11.1 | 0.27 ± 0.02 |
| Sym + (N) | -8634.5 ± 82.3 | 31.7 ± 7.9 | 0.28 ± 0.02 |
| Lat + (N) | -8649.3 ± 82.1 | 46.5 ± 11.0 | 0.27 ± 0.02 |
| P1 + P2 (N) | -8623.6 ± 81.3 | 20.8 ± 8.7 | 0.28 ± 0.02 |
| Sym + Lat + (N) | -8633.5 ± 82.3 | 30.8 ± 7.9 | 0.27 ± 0.02 |
| Sym + Lat + Sym × Lat + (N) | -8630.4 ± 82.5 | 27.7 ± 7.5 | 0.28 ± 0.02 |
| Sym + P1 + P2 + (N) | -8608.1 ± 81.5 | 5.4 ± 3.3 | 0.28 ± 0.02 |
| Lat + P1 + P2 + (N) | -8622.4 ± 81.3 | 19.6 ± 8.6 | 0.28 ± 0.02 |
| Sym + Lat + P1 + P2 + (N) | -8606.6 ± 81.5 | 3.8 ± 2.9 | 0.28 ± 0.02 |
| **Sym + Lat + Sym × Lat + P1 + P2 + (N)** | **-8602.7 ± 82.0** | **0.0 ± 0.0** | **0.28 ± 0.02** |

* Best-fit model in **bold**. Modeled effects are network (N, grouping effect), floral symmetry (Sym), latitude (Lat), and the first two principal components of the phylogenetic distance matrix (P1, P2).

**Supplementary Table 3.** Parameter estimates for the best-fit Bayesian multilevel model explaining variation in floral visitor count, $N_{visitors}$

| Effect | Estimate | 95% CI | $\hat{R}$ |
|---|---:|---:|---:|
| Std. deviation of group intercepts | 0.55 | 0.45, 0.68 | 1.00 |
| Population-wide intercept | 1.80 | 1.64, 1.96 | 1.00 |
| Symmetry | -0.26 | -0.35, -0.16 | 1.00 |
| Latitude | 0.55 | 0.38, 0.72 | 1.00 |
| Phylogenetic distance, PC1 | -0.16 | -0.20, -0.12 | 1.00 |
| Phylogenetic distance, PC2 | -0.02 | -0.06, 0.02 | 1.00 |
| Symmetry × Latitude interaction | -0.18 | -0.29, -0.07 | 1.00 |

**Supplementary Table 4.** Fit (expected likelihood predictive density, ELPD) and variation explained ($R_2$) for Bayesian multilevel models explaining variation in floral visitor sharing

| Model[*] | ELPD ± SE | ΔELPD ± SE | $R_2$ ± SE |
|---|---:|---:|---:|
| (N) | 2389.7 ± 61.5 | 7.5 ± 4.7 | 0.66 ± 0.02 |
| **Sym + (N)** | **2396.6 ± 61.7** | **0.6 ± 2.3** | **0.66 ± 0.02** |
| Lat + (N) | 2390.0 ± 61.5 | 7.2 ± 4.7 | 0.66 ± 0.02 |
| P1 + P2 (N) | 2390.4 ± 61.5 | 6.8 ± 4.3 | 0.66 ± 0.02 |
| **Sym + Lat + (N)** | **2396.7 ± 61.7** | **0.4 ± 2.2** | **0.66 ± 0.02** |
| **Sym + Lat + Sym × Lat + (N)** | **2396.9 ± 61.7** | **0.3 ± 2.1** | **0.66 ± 0.02** |
| **Sym + P1 + P2 + (N)** | **2396.7 ± 61.7** | **0.4 ± 2.2** | **0.66 ± 0.02** |
| Lat + P1 + P2 + (N) | 2390.5 ± 61.6 | 6.6 ± 4.3 | 0.66 ± 0.02 |
| **Sym + Lat + P1 + P2 + (N)** | **2397.0 ± 61.7** | **0.1 ± 1.0** | **0.66 ± 0.02** |
| **Sym + Lat + Sym × Lat + P1 + P2 + (N)** | **2397.1 ± 61.8** | **0.0 ± 0.0** | **0.66 ± 0.02** |

[*] All models with ΔELPD not significantly different from the best-fit model in **bold**. Modeled effects are network (N, grouping effect), floral symmetry (Sym), latitude (Lat), and the first two principal components of the phylogenetic distance matrix (P1, P2).

**Supplementary Table 5.** Parameter estimates for the best-fit Bayesian multilevel model explaining variation in floral visitor sharing

| Effect | Estimate | 95% CI | $\hat{R}$ |
|---|---:|---:|---:|
| Std. deviation of group intercepts | 0.69 | 0.57, 0.84 | 1.00 |
| Population-wide intercept | -1.57 | -1.76, -1.38 | 1.00 |
| Symmetry | 0.12 | 0.06, 0.18 | 1.00 |
| Phylogenetic distance, PC1 | 0.05 | 0.02, 0.08 | 1.00 |
| Phylogenetic distance, PC2 | -0.01 | -0.03, 0.02 | 1.00 |
